# From *in vitro* to *in silico*: a pipeline for generating virtual tissue simulations from real image data

**DOI:** 10.1101/2024.07.12.603259

**Authors:** Elina Nürnberg, Mario Vitacolonna, Roman Bruch, Markus Reischl, Rüdiger Rudolf, Simeon Sauer

## Abstract

3D cell culture models replicate tissue complexity, aiming to study cellular interactions and responses in a more physiologically relevant environment compared to traditional 2D cultures. However, the spherical structure of these models makes it difficult to extract meaningful data, necessitating advanced techniques for proper analysis. In silico simulations enhance research by predicting cellular behaviors and therapeutic responses, providing a powerful tool to complement experimental approaches. Despite their potential, these simulations often require advanced computational skills and significant resources, creating a barrier for many researchers.

To address these challenges, we developed an accessible pipeline using open-source software to facilitate virtual tissue simulations. Our approach employs the Cellular Potts Model, a versatile framework for simulating cellular behaviors in tissues. The simulations are constructed from real world 3D image stacks of cancer spheroids, ensuring the virtual models are rooted in experimental data. By introducing a new metric for parameter optimization, we enable the creation of realistic simulations without requiring extensive computational expertise. This pipeline benefits researchers wanting to incorporate computational biology into their methods, even if they do not possess extensive expertise in this area. By reducing the technical barriers associated with advanced computational modeling, our pipeline allows more researchers to utilize these powerful tools. Our approach aims to foster broader use of *in silico* methods in disease research, contributing to a deeper understanding of disease biology and the refinement of therapeutic interventions.

## 1 Introduction

3D cell culture models are increasingly recognized for their ability to mimic tissue complexity in terms of structure and function *in vivo*, offering transformative possibilities for biomedical research and therapeutic development (Fontoura et al., 2020; Roberto de Barros et al., 2023; Rodrigues et al., 2024). However, despite their potential, the phenotypic quantitative 3D image analysis from spheroid wholemount microscopy data sets remains a significant challenge, as accurate visualization and characterization of cells throughout these structures are hampered by technical and analytical limitations (Tang et al., 2023). There is also a growing interest in complementing these experiments with *in silico* simulations to enhance research efficiency and outcomes, including aspects like cost-effectiveness, speed, predictive power, and the ability to integrate diverse data sources. These computational models provide a powerful tool for exploring cellular behavior and interactions within 3D environments without the constraints of traditional experimental techniques (Jean-Quartier et al., 2018; Berghoff et al., 2020; Cortesi et al., 2021).

*In silico* simulations of tumor spheroids have become an important tool in cancer research, offering complementary capabilities to traditional experimental methods for studying tumor growth and treatment response. By integrating complex biological simulations of tumor dynamics under various conditions, these computational models provide insights often difficult to obtain through *in vitro* or *in vivo* studies alone. By accurately capturing the interactions between cell proliferation, nutrient diffusion, and tumor microenvironment, *in silico* models can predict tumor behavior and assess the response to therapeutic interventions and their effectiveness (Kam et al., 2012; Amereh et al., 2023; Hickey et al., 2024). These studies, which utilize *in silico* modeling of cancer spheroids, highlight the critical role of *in silico* simulation in advancing our understanding of cancer biology and the development of personalized medicine strategies. Cell-based computational models include lattice and off-lattice methods, each with distinct advantages and limitations. Lattice methods, such as Lattice-Gas Cellular Automata (LGCA), Cellular Automata (CA), and Cellular Potts Model (CPM), vary in resolution and computational efficiency, with CPM offering the highest morphological detail. Off-lattice methods, including Center-Based Models (CBM), Subcellular Element Models (SEM) and Boundary-Based Models (Vertex and Front Tracking), provide higher resolution and flexibility for simulating detailed cell shapes and interactions but are more complex and computationally intensive (Metzcar et al., 2019). Therefore, these simulations are challenging to implement and typically require access to computing clusters or high-performance PCs. Moreover, they demand specialized skills in bioinformatics or coding—expertise that may not be readily available in every laboratory. Consequently, not all research facilities have the immediate computational power or specialized knowledge to leverage these advanced technologies.

To facilitate the generation of virtual tissue simulations, we have developed a new pipeline, which relies exclusively on open-source software and requires minimal coding skills. The process starts with obtaining 3D image stacks of optically cleared cancer spheroids via confocal microscopy, followed by 3D image segmentation at the single-cell level. The segmentation result is then imported into CompuCell3D (CC3D) (Graner and Glazier, 1992; Glazier and Graner, 1993), a simulation environment for virtual tissue modeling based on the CPM. To steer the simulation towards more realistic simulation results, we defined a new metric based on the statistical distribution of morphological features, and used it during parameter optimization to evaluate simulation outcomes. One specific use of this pipeline was already demonstrated by using simulated cancer spheroids as input to machine learning approach to generate synthetic images as training data for image segmentation algorithms (Bruch et al., 2024, submitted manuscript).

## 2 Materials and Methods

### 2.1 Cell culture

For the generation of mono-culture spheroids, the colorectal adenocarcinoma cell line HT-29 (ATTC) was cultured as previously described (Nürnberg et al., 2020). Briefly, cells were cultured in McCoy’s 5A medium (Capricorn) supplemented with 10% fetal bovine serum (FBS, Capricorn) and 1% penicillin/streptomycin (Sigma-Aldrich). For spheroid generation, cells were detached using Trypsin/EDTA and seeded onto 96-well ultra-low attachment (ULA) U-bottom plates (Corning) at a concentration of 500 cells per well. Spheroids formed by a self-driven mechanism over 3 days. Cells were maintained in a humidified incubator at 37°C and 5% CO2, repeatedly authenticated by phenotypic analysis, and regularly tested for mycoplasma.

### 2.2 Immunofluorescence and optical clearing

Spheroids were fixed and fluorescently labeled as previously reported (Nürnberg et al., 2020). If not stated otherwise, all steps were carried out at room temperature (RT) under gentle agitation of samples. Briefly, spheroids were transferred into 200 μL PCR tubes, washed twice with phosphate-buffered saline (PBS), and subsequently fixed with 4 % paraformaldehyde (PFA, Carl Roth) for 1 h at 37 °C, followed by washing twice with PBS containing 1% FBS, for 5 min each and permeabilization in 2 % Triton X-100 diluted in PBS for 5-10 min. Then, spheroids were quenched with 0.5 M glycine (Carl Roth) in PBS for 1 h at 37°C and subsequently incubated in penetration buffer (0.2 % Triton X-100, 0.3 M glycine, 20 % dimethyl sulfoxide (DMSO, all Carl Roth) in PBS) for 30 min. Samples were then washed twice with PBS/1 % FBS, followed by incubation in blocking buffer (0.2 % Triton X-100, 1 % bovine serum albumin (BSA, Carl Roth), 10 % DMSO in PBS) for 2 h at 37 °C. After blocking, samples were incubated with primary antibody overnight (ON) at 37°C with gentle shaking. Primary anti-KI67 antibody (Merck, rabbit polyclonal antibody) was diluted 1:300 in antibody buffer (0.2 % Tween 20, 10 μg/mL heparin (both Sigma-Aldrich), 1 % BSA, 5 % DMSO in PBS). Following primary antibody incubation, samples were washed 5 times for 5 min each in washing buffer (0.2% Tween 20, 10 μg/mL heparin, 1 % BSA) and stained with membrane dye, secondary antibody, and nuclear dye ON at 37°C with gentle shaking and protected from light. Corresponding secondary antibody and dyes were diluted in antibody buffer with the following concentrations: donkey anti-rabbit AlexaFluor488, 1:800 (Invitrogen); DAPI, 1:1000 (Sigma-Aldrich); SiR-actin, 1:1000 (Spirochrome). Samples were washed subsequently 5 times for 5 min in washing buffer with gentle shaking followed by sample immersion in FUnGI clearing agent (50 % glycerol (vol/vol), 2.5 M fructose, 2.5 M urea, 10.6 mM Tris Base, 1 mM EDTA) for several hours until sample transparency was sufficient (Rios et al., 2019).

### 2.3 Microscopy

All images were acquired with a Leica TCS SP8 confocal microscope (Leica Microsystems CMS, Mannheim) equipped with HC PL APO 20×/0.75 IMM CORR objective, 405 nm, 488 nm, and 633 nm lasers. Image stacks were acquired at a resolution of 1024×1024 pixels, a z-step size of 2 µm (voxel size xyz: 0.5682×0.5682×2.0 µm^3^), and a line average of two. Settings for laser intensity and gain were chosen in a sample-dependent manner such that overexposure of pixels was avoided.

### 2.4 Image processing and segmentation

All image stacks were pre-processed in FIJI by applying a manual background correction, which involves increasing the minimum grey value of an image in an image-dependent manner, and 3D Gaussian smoothing (σ = 0.4 px). Segmentation of the cell membrane staining was performed with Cellpose 2.0 and a custom model based on the pre-trained cyto2 model (Stringer et al., 2021; Pachitariu and Stringer, 2022). The nuclear channel was included in the training process of the membrane segmentation to improve results. Model training was performed as human-in-the-loop training by applying the pre-trained cyto2 model to a single-plane image of the spheroid, followed by manual correction of label masks. The training was then performed based on the corrected label masks with a learning rate of 0.1, a weight decay of 10^−5^, and 300 epochs (Cellpose default settings), and the trained model was applied to the next single-plane image. After processing 20 single-plane images, the quality of the membrane segmentation reached a level where no further major manual adaptations were necessary. The trained segmentation model was then applied to entire image stacks of optically cleared HT-29 spheroids with a flow threshold of 0.4, a cell probability threshold of 2.0, and a stitch threshold of 0.45 (Supplementary Figure S1). Segmentation quality was assessed via detection and segmentation accuracy measured as described in (Matula et al., 2015). The outcome of the segmentation process is a new image, referred to as a label mask, where each individual cell is represented by a unique identifier label.

### 2.5 Feature extraction and statistical analysis

A custom-made Python script was generated for feature extraction and quantitative analysis of 3D label masks. It makes use of the *NumPy* and *sci-kit-image* libraries (van der Walt et al., 2014; Harris et al., 2020).

#### Pre-processing of label masks

To reduce remaining segmentation errors, small labels were eliminated by removing objects with a volume smaller than 5 voxels, followed by closing and opening operations. Labels present in only one or two planes were merged with adjacent labels, provided these adjacent labels also spanned only one or two planes, as they likely originated from segmentation errors. In cases where adjacent labels did not meet this criterion, the small labels were removed from the image to minimize residual segmentation errors.

To ensure an isotropic voxel size, which is, later on, necessary for CC3D simulations, image upsampling (with nearest-neighbor interpolation) was applied in z to match the x-y resolution. The data presented in this manuscript is derived from an original image stack with a size of 517×517×136 voxel (xyz), which was upsampled to a final resolution of 517x517x479 voxel (xyz).

#### Morphological feature extraction

The following morphological properties of individual cells were extracted using the *regionprops* function from the *scikit-image* library: volume, minor and major axis length, and eccentricity. Additionally, the surface area was approximated via the sum of border voxels, as well as the volume-to-surface (V/A) ratio and the sphericity of cells. These morphological features are later used to calculate the deviation of the simulation from the real-world images, and to determine model parameters that minimize this deviation.

### 2.6 Simulation and parameter optimization

Biophysical simulations were carried out with CC3D (Version 4.3.2, Revision 0). This advanced simulation environment for modeling of complex biological phenomena through biophysical principles is based on the Cellular Potts Model (CPM). In the CPM approach, each pixel (2D) or voxel (3D) is assigned to a specific cell or to the extracellular medium. Then, the total free energy of the system is expressed as a Hamiltonian function *H*, and the Monte Carlo algorithm is used to evolve the initial cellular configuration into its thermodynamic equilibrium (Graner and Glazier, 1992; Glazier and Graner, 1993; Swat et al., 2012). The Hamiltonian can incorporate various biophysical interactions such as cell adhesion, volume constraints, and surface tension according to the differential adhesion hypothesis (DAH) (Steinberg, 2007). In this way CC3D can be used to simulate dynamic cellular processes over time, making it particularly well-suited for studying developmental biology, tissue engineering, and cancer research.

#### Transfer of segmented label mask to CC3D

Usually, simulations based on the CPM are initialized with an artificial configuration, such as stacked cubes. From there, a sophisticated model and numerous steps in the Monte Carlo simulation are required in order to converge to a biologically realistic configuration. For this reason, in this work real-world 3D confocal image stacks and the corresponding image segmentation are used as initial configuration in the simulation. By starting the simulation from a realistic configuration, also less sophisticated models can lead to realistic short-term dynamics of the system, with less computation effort.

For the transition from real image data to an *in silico* simulation with CC3D, a custom Python script was used to generate the necessary Potts Initial File (PIF), which defines the starting configuration of the simulation. Starting from the pre-processed label image, the PIF consists of one line for every voxel that is occupied by a cell, using the following syntax:

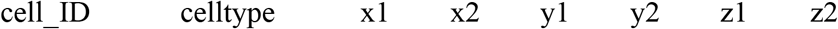

Here, cell_ID is the unique identifier for a cell and corresponds to the label ID assigned during image segmentation, and celltype is a string that corresponds to a cell type. In the present work, we only deal with a single cell type. In general, cell types are defined in the .xml file of the simulation. The remaining elements represent a range of coordinates in x-, y-, and z-direction, encompassing the voxels occupied by a cell with a given cell ID and type. In the CC3D syntax, x1, y1, and z1, along with x2, y2, and z2, represent the coordinates of the corners of a bounding box that defines the spatial boundaries of a cell, with the indices 1 and 2 indicating the opposing corners in each plane. Since each cell, in our case, needs to be represented by individual voxels rather than a bounding box, x1 and x2, as well as the corresponding y and z coordinates, are identical, so each line in the file represents a single voxel.

#### CC3D simulation

Initial simulations were generated using CC3D’s simulation wizard (implemented in Twedit++). A detailed list of selected properties and plugins can be found in Supplementary Table S2. For a detailed explanation on the software, we recommend and refer to the CC3D documentation (Swat et al., 2012).

Cell types were specified as defined during PIF file generation, as well as an additional cell type termed “Wall”, which serves as a barrier between cells and the border of the lattice to prevent cells from sticking to the boundaries.

The central component of the CPM is the system Hamiltonian ***H*** (Eq. 1), which models the overall free energy of the cell cluster. Here, we consider the adhesion energy between adjacent cells and the surface free energy of cells in contact with the surrounding medium. Additionally, the Hamiltonian includes penalty terms to account for any deviation of cell i’s volume V_i_ and surface area A_i_ from their respective target values:

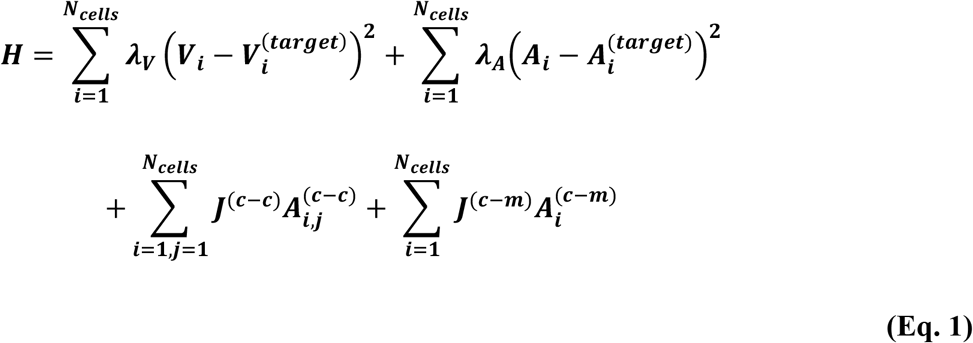

In this context, λ_V_ and λ_A_ represent penalty parameters that dictate the intensity of the volume and area constraints, respectively. The term 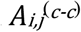 denotes the contact area between cell *i* and cell *j*, while 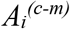 refers to the contact area of cell *i* with the surrounding medium. Additionally, *j*^*(c-c)*^ and J^(c-m)^ are the corresponding surface tension parameters. Individual target values for volume 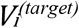 and area 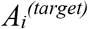 were specified for each cell. At the start of the simulation, each cell is set to be already at its target volume and surface, except for cells smaller than 15% of the average volume or larger than the average plus two standard deviations. For these cells, a set of target volumes and surfaces is randomly assigned, sampled directly from cells that fall within the specified range of properly segmented cells. A scatter matrix plot illustrating extracted morphological features for both correctly segmented cells and those assigned a new volume and surface can be found in Supplementary Figure S3.

All simulations were carried out on a workstation equipped with a 16-core processor, 128GB of RAM, and an 8GB GPU.

#### Simulation output

Several output quantities were monitored during the simulation to track the progression and morphological changes over time. In addition to the average cell volume and surface, the current simulation state was exported and saved as a .tiff file at defined Monte Carlo Steps (MCS) for further processing and analysis. Furthermore, custom functions were implemented to track and export morphological features of each simulated cell in defined intervals for subsequent analyses. Multiple similarity parameters were calculated and exported at defined intervals to assess cellular morphology changes over time. In more detail, for each morphological parameter, the first Wasserstein distance *W* (Eq. 2, Ramdas et al., 2015) between the frequency distributions at the start and a particular MCS is calculated, as well as the intersection over Union (IoU, Eq. 3) of a cell at a particular MCS compared with its starting position, averaged over all cells:

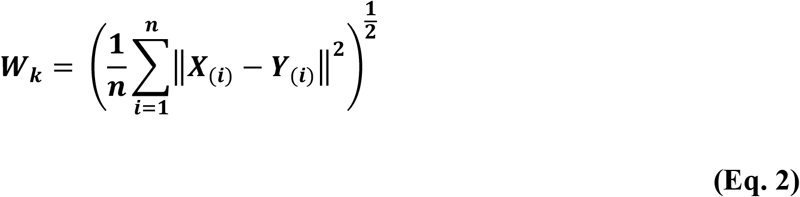

Here, *X*_*(i)*_ and *Y*_*(i)*_ are the ordered values of the morphological feature *k* at the start and a particular MCS.

In our context, *A*_*i*_ is the volume of a cell *i* at the start of the simulation, and *B*_*i*_ is the volume of the same cell at a particular MCS.

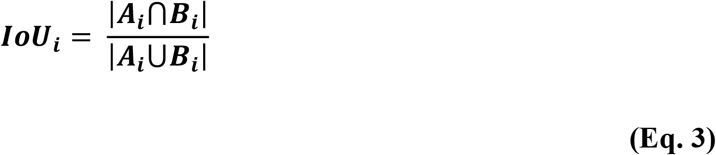

In our context, A_i_ is the volume of a cell *i* at the start of the simulation, and B_i_ is the volume of the same cell at a particular MCS.

#### Parameter optimization

To achieve realistic simulation results, an optimization of model parameters had to be carried out. These parameters are the contact energy between cells and the medium (*J*^*Cell-Medium*^), contact energy between neighboring cells (*J*^*Cell-Cell*^), and the energy constraints *λ*_*Volume*_ and *λ*_*Surface*_, which act as penalty parameters in case of deviations from the specified target cell volume and surface area. Since the target volume and surface of cells were taken from the real-world image, these parameters were not included in the optimization. Parameter optimizations involved scanning different combinations of the mentioned parameters and were performed on a small image patch with a size of 125x125x178 pixels (xyz) to reduce computation time. For this, the preceding steps of the pipeline were conducted as described above. Additionally, incomplete cells at the edges of the image caused by image cropping were manually removed after image segmentation before generating the PIF file.” After setting up the basic simulation, the following parameters were selected to be altered between individual scans (Table 1):

**Table 1.** Parameters included in parameter optimization scans.

| PARAMETER | VALUES |
| --- | --- |
| $J_{Cell-Medium}$ | 10.0 / 55.0 / 100.0 J/pix<sup>2</sup> |
| $J_{Cell-Medium}$ | 2.0 / 4.0 / 6.0 / 8.0 / 10.0 J/pix<sup>2</sup> |
| $\lambda_{Volume}$ | 0.001 / 2.0 / 4.0 / 6.0 / 8.0 / 10.0 J/pix<sup>6</sup> |
| $\lambda_{Surface}$ | 0.001 / 2.0 / 4.0 / 6.0 / 8.0 / 10.0 J/pix<sup>4</sup> |

The remaining parameters were kept at constant values and can be found in Supplementary Table S2. The overall target of the parameter optimization was to identify model parameters that maintain the overall shape of the distributions of morphological features across MCSs, while preventing cell fragmentation and ensuring sufficient variation of the cellular shape from its initial form. An attempt to condense these goals in a target function for parameter optimization is a metric WIP (Wasserstein-IoU product) that combines the mean Wasserstein distances between the start and end of the simulation and the mean IoU between cells:

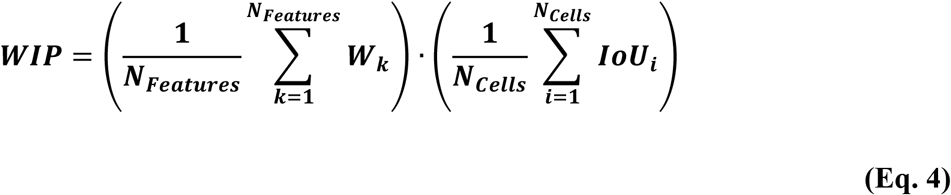

Here, *N*_*Features*_ represents the total number of the morphological features, *k* is the index for a specific feature, *N*_*Cells*_ denotes the total number of cells, and *i* is the index for a specific cell.

The first factor of WIP (Eq. 4), the mean Wasserstein distance, ensures that cells do not significantly deviate from their original morphological features. However, solely relying on the Wasserstein distance can lead to parameter sets with high volume/area penalty parameters (λ_Volume_ and λ_Surface_) being favored, since in this case, the cells are frozen due to high energy constraints. The second factor, the mean IoU, is used to ascertain that cells, while maintaining the similarity of the ensemble, still exhibit a certain degree of shape variability on the level of individual cells. Otherwise, static simulations would be favored.

## 3 Results

### 3.1 Segmentation of optically cleared HT-29

High-resolution confocal microscopy on optically cleared spheroid samples revealed detailed morphological characteristics of HT-29 cancer spheroids at the cellular level. The use of optical clearing during fluorescence labeling allowed for high-quality 3D image stacks throughout the entire sample, mitigating significant signal loss towards the center due to light scattering. This approach enabled the acquisition of detailed single optical sections from a 3D confocal image stack, providing comprehensive insights into the cellular shape and distribution of fluorescence signals across the spheroids (Figure 1A). Since downstream analyses require high-quality fluorescence images of both, cell nuclei and membrane, DAPI was used to label cell nuclei and cell membranes were visualized using SiR-Actin, marking the F-actin cortex underneath the cell membrane. Subsequently, 20 randomly selected single optical sections from acquired image stacks were used for training a custom Cellpose 2.0 model to segment cell boundaries across the sample (Figure 1B). This specific number of sections was selected to establish a balanced approach between time investment into training and the accuracy of the segmentation. In total, the time needed for training the model and manually correcting 20 single optical sections was 3 hours. However, deploying this trained model using the 3D functionality of the Cellpose software led to imprecise membrane segmentation. Therefore, the model was instead utilized in stitching mode, which performs slice-wise 2D segmentation initially. In a subsequent step, it merges overlapping regions between adjacent labels in the z-direction to determine whether a label corresponds to the same cell or a different one. Despite achieving substantially better results, minor segmentation errors were still present and accepted in the label masks to balance time investment in the training process and segmentation accuracy (Figure 1C). The overall segmentation quality was assessed using the detection (DET) and segmentation (SEG) scores against two manually segmented ground truth sets, resulting in scores of 0.73 and 0.46, respectively (Supplementary Figure S1).

**Figure 1.**
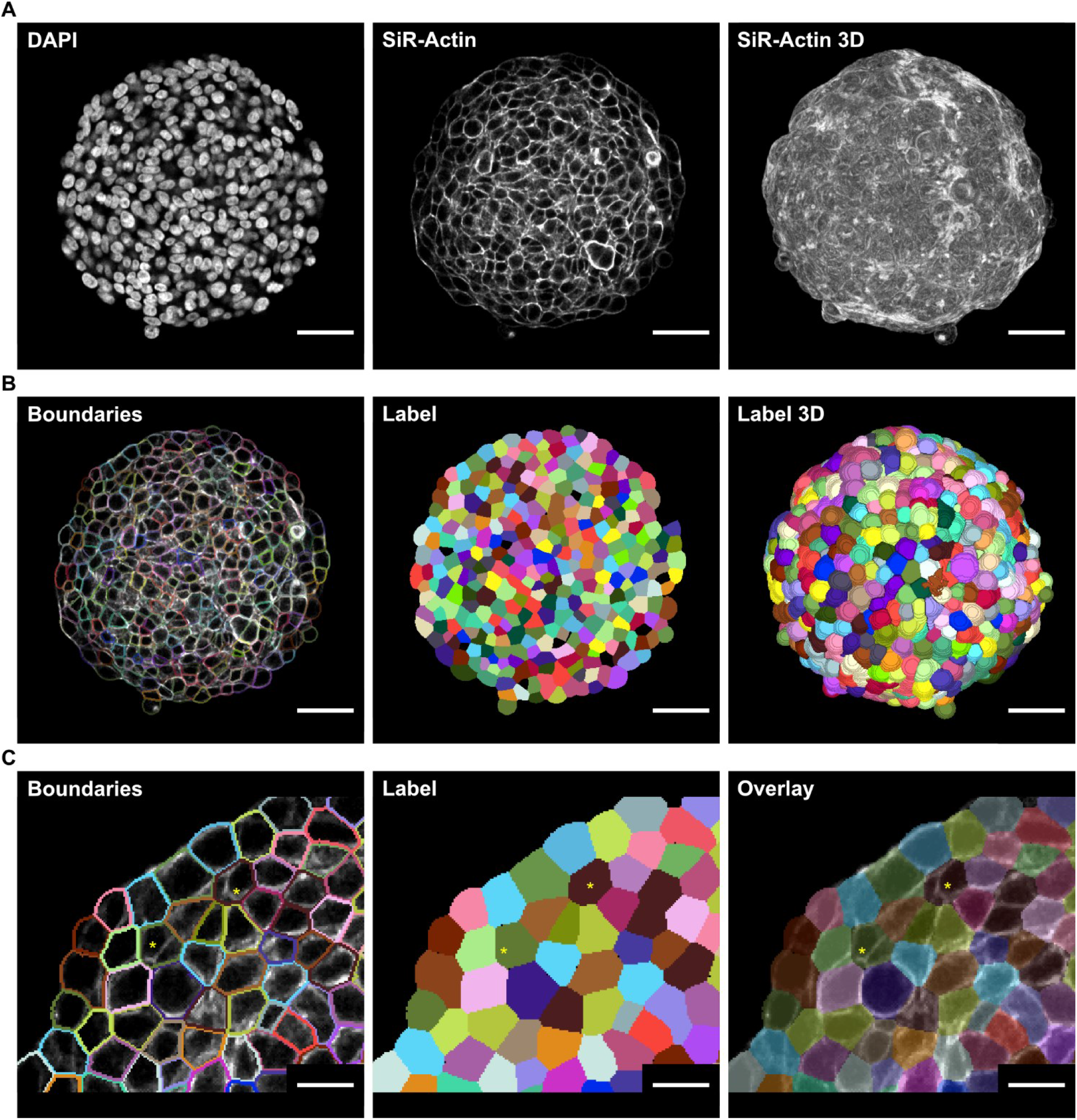
Optical clearing enhances 3D microscopy of HT-29 spheroids for membrane segmentation. Spheroids of HT29 cells were generated by seeding 500 cells/ well onto ULA multi-well plates and then cultured for three days. Following PFA fixation, fluorescence labeling with DAPI and SiR-actin, and optical clearing, samples were imaged with wholemount confocal microscopy. (A) Raw real-world images of a representative spheroid showing DAPI (left) and SiR-actin signals of a single optical section through the central portion (middle) or a maximum-z projection of the entire spheroid (right) (B) Cell boundaries (left) and label masks of cells from the optical section shown in A (middle), and 3D labels of the entire segmented spheroid (right) after Cellpose training. The number of sections used for training was chosen to balance the time investment required and acceptance of minor segmentation errors. (C) Detail of segmentation from B depicting cell boundaries over the raw image (left), cell labels (middle), and translucent cell labels over the raw image (right) Segmentation errors are indicated by asterisks. Scale bars: 50 µm (A, B), 20 µm (C).

The trained model was applied to entire image stacks of spheroids to segment cell boundaries and generate label masks on a single-cell level (Figure 1B, right panel). Cell labels were post-processed, as described in Section 2.5, to filter out segmentation errors and to generate the configuration file for the CC3D simulation. All label masks were analyzed further to extract morphological features, such as cell volume, surface, major and minor axis length, eccentricity, and sphericity.

### 3.2 Generation of whole spheroid simulations from real image data

The virtual tissue simulation environment CC3D was then used to simulate an entire spheroid based on previously acquired real-world image data of HT-29 spheroids as the basis of the cellular layout in a lattice with the original image stack dimensions (Figure 2, MCS0). Figure 3 shows a simulation that was conducted using optimized model parameters (section 3.3). Visual examination of different stages during the simulation revealed that with increasing simulation progress, the cells increasingly lost their initial irregular shapes and converged into a less cluttered and more harmonious configuration. The cell boundaries are initially jagged and complex but smooth out over time, and the shape of the cells becomes more regular and rounded, thereby reducing the boundary complexity. (Figure 2, lower row).

**Figure 2.**
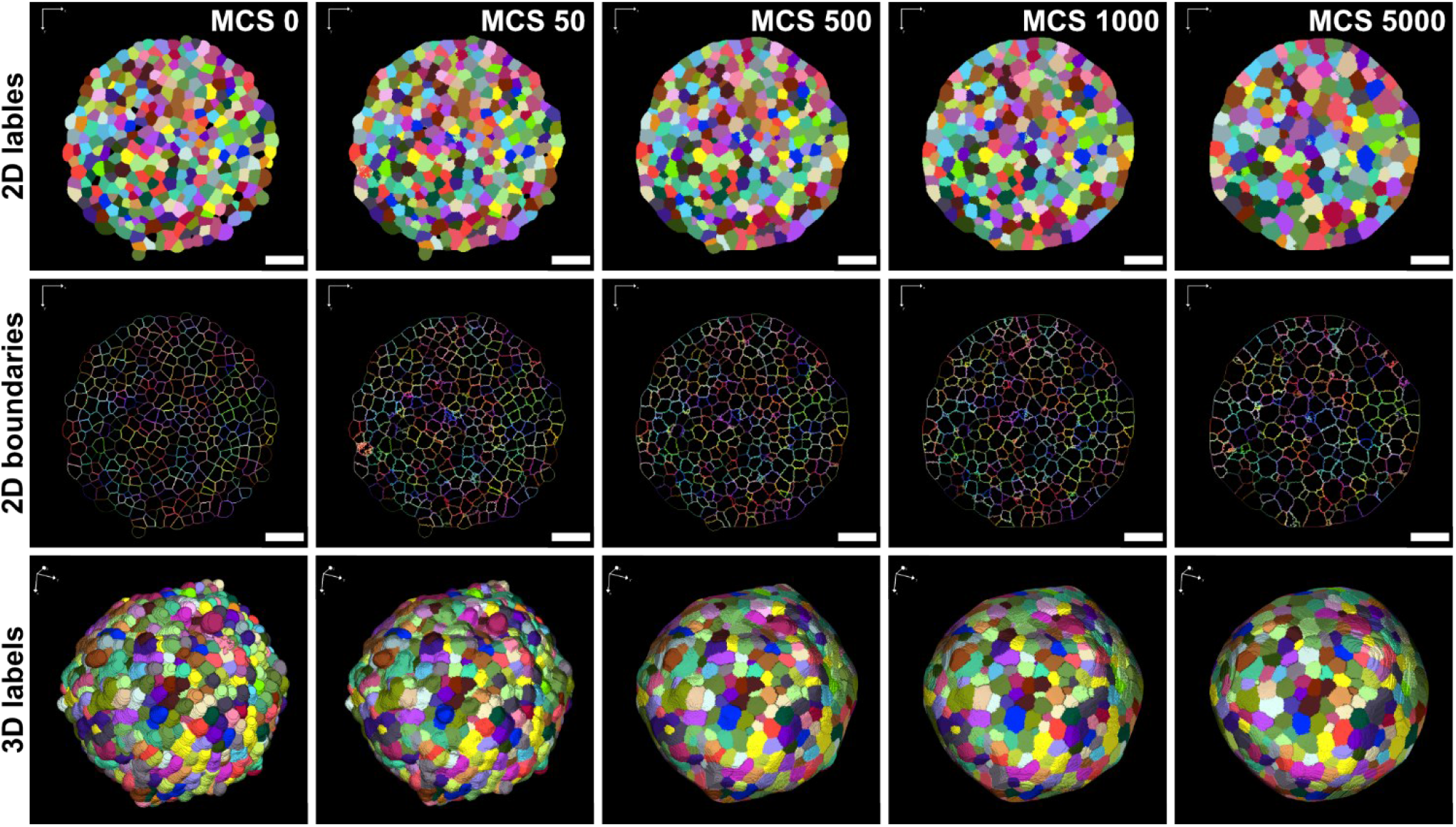
Generation of virtual 3D simulations from real-world image data of HT-29 spheroids. Label masks of segmented HT-29 spheroids were used to create the starting configuration of cells for a CC3D simulation (MCS 0). Figure shows the label and boundaries of individual cells in 2D (upper and middle panels) and 3D label masks (lower panels) at distinct time points during the simulation. Number of Monte Carlo Simulations (MCS) is indicated in the upper right corners of upper panels and applies also all corresponding panels in the same column. Parameters were set as follows: λ_V_ = 10.0 J/pix^6^, λ_A_ = 0.001 J/pix^4^, J^(c-c)^ = 2.0 J/pix^2^, J^(c-m)^ = 55.0 J/pix^2^. Scale bars: 75 µm

**Figure 3.**
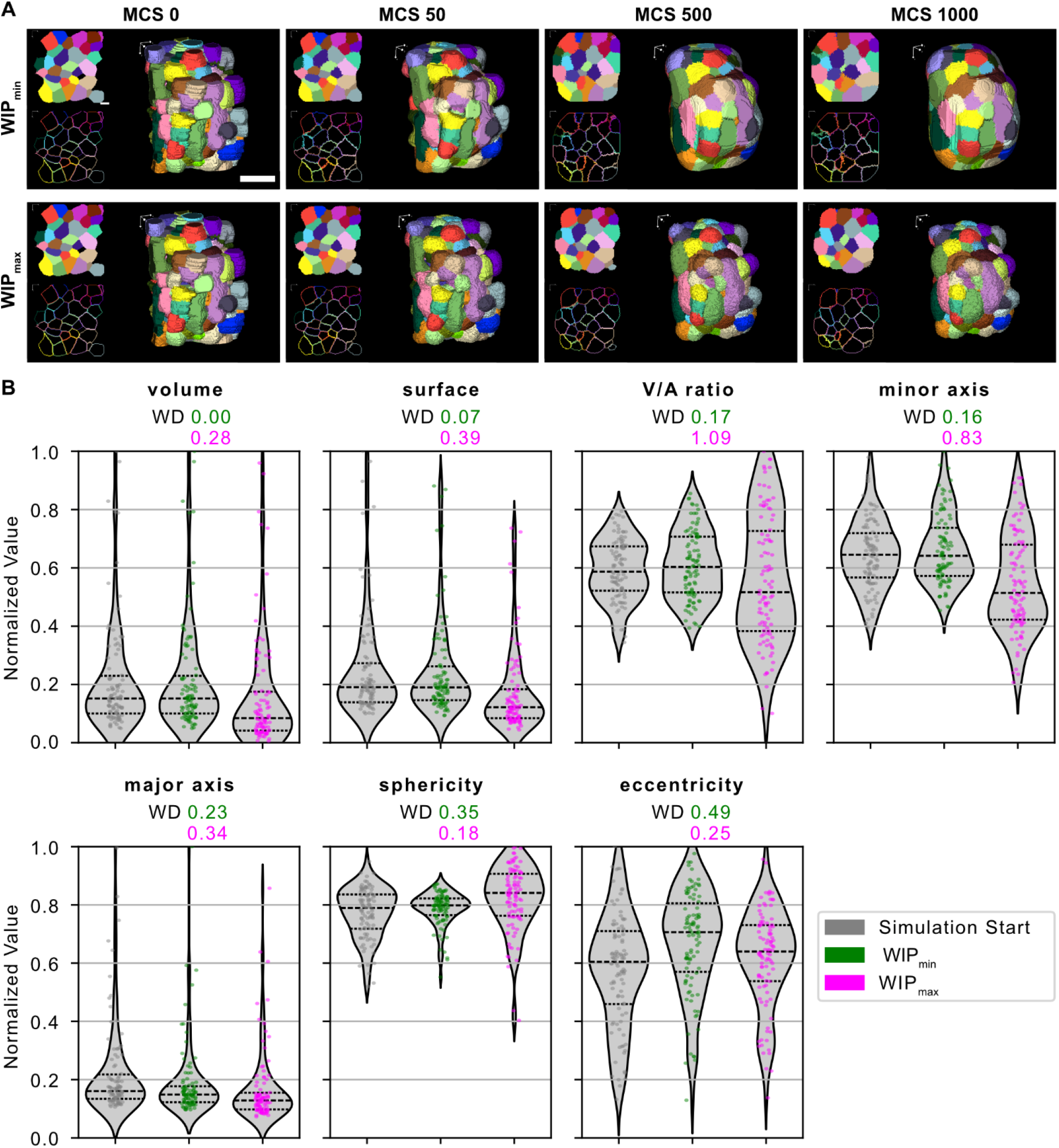
Model parameter optimization. Different combinations of values for volume constraint λ_V_, surface constraint λ_A_, and the contact energies J^(c-c)^, and J^(c-m)^ were tested on a small subset of manually segmented cells of an HT-29 spheroid and analyzed for minimization of the metric *WIP* (Eq. 4). (A) depicts exemplary 3D projections of simulated cells at distinct MCS for a simulation with parameters resulting in the minimum (upper panel) and maximum (lower panel) value of *m*. (B) shows violin plots of the morphological feature distributions at the simulation start and end of the simulations, with the smallest (green) and largest (magenta) values for the metric WIP. Dashed lines correspond to the distribution’s median and the first and third quartiles. The respective WD between the start and end of a simulation is shown above each plot. Scale bar: 50 µm. Parameters: WIP_min_: λ_V_ = 10.0 J/pix^6^, λ_A_ = 0.001 J/pix^4^, J^(c-c)^ = 2.0 J/pix^2^, J^(c-m)^ = 55.0 J/pix^2^; WIP_max_: λ_V_ = 0.001 J/pix^6^, λ_A_ = 10.0 J/pix^4^, J^(c-c)^ = 10.0 J/pix^2^, J^(c-m)^ = 10.0 J/pix^2^)

### 3.3 Parameter optimization based on the distribution of cellular morphological features

Since the default parameters of CC3D appeared to be unsuitable for maintaining cellular morphology, a parameter optimization was conducted to adjust model parameters that affect cellular morphology towards more realistic simulation outcomes. The results were ranked according to the metric WIP (Eq. 4), which ensures that individual cells undergo shape changes during the simulation while the overall feature distribution remains stable. Parameter optimizations were performed on a small, manually segmented subset of a larger image stack. The results indicated that the optimal performance, as measured by the lowest value of metric WIP, was achieved under specific conditions: a low surface constraint (λ_A_) of 0.001 J/pix^4^ and a larger volume constraint (λ_V_) of 10.0 J/pix^6^ (Figure 3). Analyzing the calculated Wasserstein distances (WDs) for various morphological features showed that an elevated volume constraint resulted in a highly consistent distribution of cellular volumes between the start and end of a simulation. To a lower degree, a similar consistency was observed regarding the distributions of cell surface area, the volume-to-surface area ratio (V/A), and the lengths of the major and minor axes across the simulated spheroid. Despite the parameter set achieving the lowest metric WIP, it is important to note that the shape descriptors—sphericity and eccentricity—exhibited larger WDs than those obtained from simulations where WIP was large (Figure 3B). This is reflected in the simulated cells displaying a more rounded shape and less morphological change over the course of the simulation (Figure 3A, lower panel), as compared to the overall smoothing of the surface and shape changes observed for smaller values of WIP (Figure 3A, upper panel).

## 4 Discussion

We have generated a new pipeline to facilitate *in silico* simulations in 3D cell culture research. This pipeline uses solely open-source software and requires minimal coding expertise. The process begins with acquiring 3D image stacks of optically cleared cancer spheroids through confocal microscopy, followed by segmentation of the entire spheroid on a single cell level using Cellpose 2.0. The resulting label masks are then transferred to CC3D, a simulation environment for virtual tissue simulations.

### Whole mount segmentation of HT-29 spheroids

The initial phase of setting up an *in silico* simulation of HT-29 spheroids involved segmenting the membranes of all cells within the spheroid. The segmentation model used in this research was trained on a limited number of optical sections to balance time expenditure and segmentation accuracy. Despite achieving good segmentation results for single optical sections, some errors persisted, particularly at the top and bottom of cells. This was mostly due to the difficulty in distinguishing between individual cells in these regions, even with manual segmentation. To address this issue, we suggest adjusting the microscope parameters during image acquisition by increasing the resolution and reducing the step size. This is especially important for 3D segmentation in stitching mode, as missing information between single optical planes can substantially affect the process.

### Generation of virtual tissue simulations from real-world image data

Using the results of image segmentation as a starting point for *in silico* simulations allows for the setup of initial configurations that closely resemble the real *in vitro* system, thus bypassing the often time-consuming and complex process of manually developing and validating a simulation model that starts from artificial building blocks and develops into a real cell configuration. Furthermore, this method can be further developed to simulate biological systems by incorporating additional biological processes. For example, in many *in vitro* studies using spheroid models, a fundamental interest is the proliferation rate in 3D cell cultures, which often differs from traditional 2D cell culture models.

The present method includes only the morphological characteristics of cells and does not incorporate additional biological phenomena. However, it can be easily extended. For example, many experiments involve visualizing proliferation markers, such as KI67, at distinct time points during cultivation. To model cell proliferation, researchers can perform additional segmentation of KI67-positive cells or use statistical methods to cluster cells by their KI67 signal intensity. This approach allows for incorporating additional cell types into the simulation, enabling the modeling of cell proliferation by implementing chemical nutrient fields and using the ratio of proliferating cells as an optimization parameter.

Furthermore, the software used in this pipeline is open-source and can be run on modern workstations. However, as the number of the simulated voxels increases, which was already substantial in this study (478x464×479 voxel), computational limits are quickly reached, necessitating more powerful computing resources. In our case, whole spheroid simulations required a computation time of 62 hours for 5000 MCS.

### Parameter optimization

For parameter optimization, performed on a small, manually segmented subsection of a spheroid, we focused on the metric WIP that measures the consistency of the distribution of morphological characteristics of cells during the simulation with the *in vitro* experiment, whilst enforcing sufficient dynamic changes during the simulation. By comparing feature distributions at the start of the simulation, identical to the segmentation result, and at the end of the simulation, the mean WD ensured a comparable distribution of features throughout the simulation. The average IoU prevented the optimization of simulation parameters towards static or “frozen” cells. In our case, the focus was on the statistical similarity of feature distributions, making the minimization of this metric our primary goal. However, this focus led to visible differences in the appearance of cells, particularly in the border regions of the spheroid. This suggests that the model does not adequately reflect the necessary contact energies to maintain the specific shapes of cells in peripheral regions. This outcome is not surprising, as it is well known that cells in a 3D aggregate exhibit different biological behavior depending on their location. For example, proliferating cells tend to concentrate toward the border region due to a higher nutrient supply than cells in the spheroid center. To better reflect this behavior in *in silico* simulations, a potential solution is the introduction of additional cell types into the simulation, each with its own set of model parameters.

In conclusion, the presented pipeline offers a robust and accessible approach for simulating 3D cellular aggregates, which can be performed on a single workstation using exclusively open-source software. This method requires limited coding skills, making it accessible to a broader range of researchers. The primary input for the method is real-world image data, allowing for straightforward implementation and analysis. The model can be further enhanced by incorporating more detailed biological information, such as additional cell types via specific markers like KI67 for proliferation and spatial factors like a cell’s distance from the surface. This flexibility ensures that the method can be adapted to reflect various biological processes more accurately, enhancing the relevance and applicability of *in silico* simulations in biological research.

## 6 Author Contributions

Conceptualization, E.N. and S.S.; methodology, E.N. and S.S.; software, E.N., M.V. and R.B.; validation, E.N. and S.S.; formal analysis, E.N.; investigation, E.N., M.V.; resources, S.S., M.R. and R.R.; data curation, E.N.; writing—original draft preparation, E.N. and S.S.; writing—review and editing, M.V., R.B., M.R., R.R.; visualization, E.N.; supervision, S.S., M.R., and R.R.; project administration, S.S., R.R. and M.R.; funding acquisition, S.S., M.R., R.R.. All authors have read and agreed to the published version of the manuscript.

## 8 Acknowledgments

We acknowledge Felix Romer for his valuable assistance and contributions to this project.

## 9 Data Availability Statement

Datasets are available on request: The authors will make the raw data supporting this article’s conclusions available.

