## Supplementary Files for "From *in vitro* to *in silico*: a pipeline for generating virtual tissue simulations from real image data"

### *Supplementary Material*

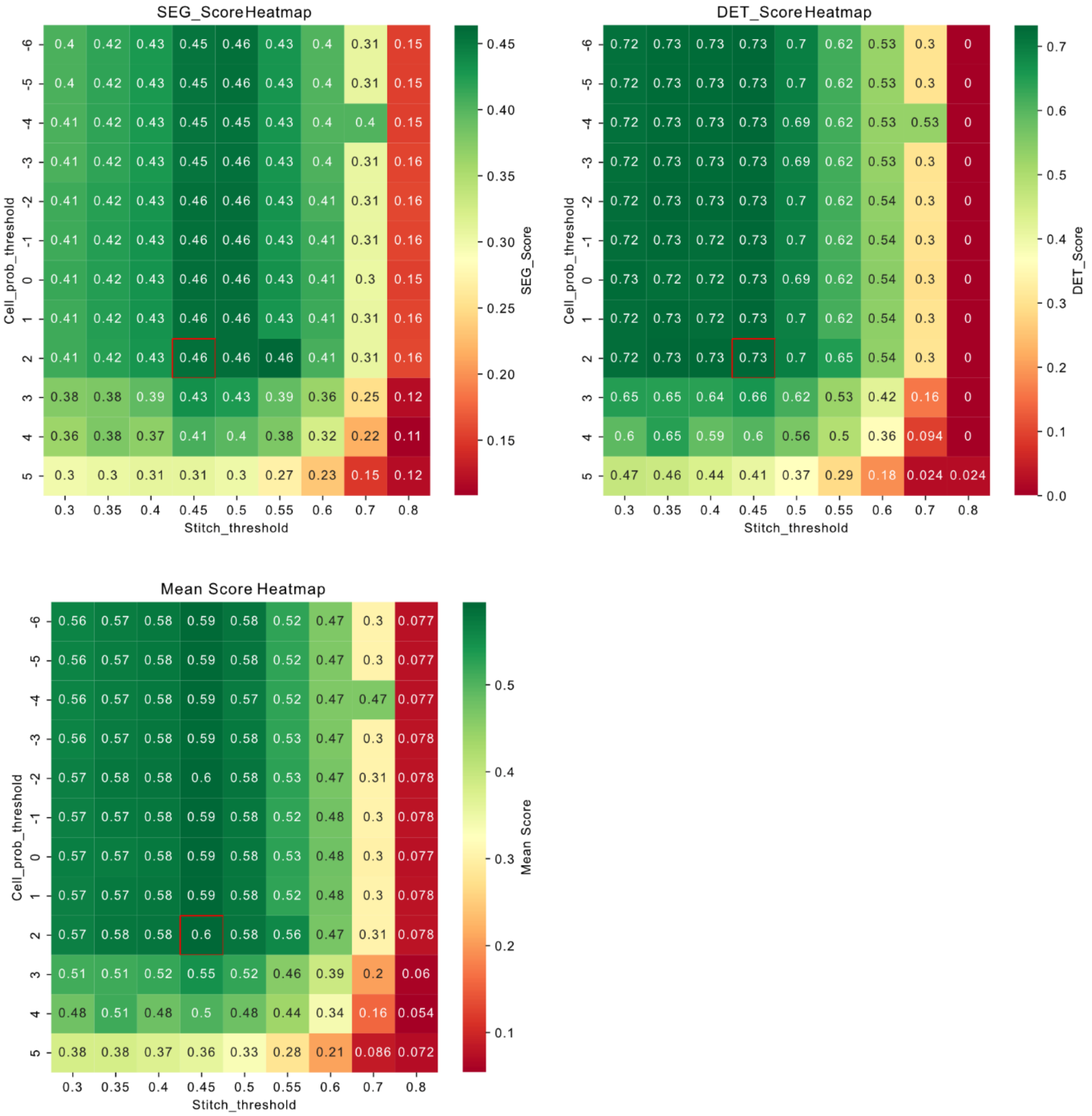


**S 1 Evaluation of segmentation quality for different threshold parameters in Cellpose 2.0.** Performance of the trained segmentation model in Cellpose 2.0 was evaluated for different cell probability and threshold parameters by using the segmentation (SEG_Score, upper left) and detection score (DET_Score, upper right) as described in the Cell Tracking Challenge (Matula et al., 2015). The final threshold parameters were determined by calculating the mean of both scores (lower left), as indicated by the red rectangles.

**S 2 Summary of simulation properties used to generate basic CC3D simulations**

| | |
|---|---|
| **GENERAL SIMULATION PROPERTIES** | |
| **LATTICE DIMENSION** | Dimensions of the image stack after upsampling |
| **BOUNDARY CONDITIONS** | No flux in x, y, and z |
| **LATTICE TYPE** | Square |
| **AVERAGE MEMBRANE FLUCTUATIONS** | 30 |
| **PIXEL COPY RANGE** | 3 |
| **NUMBER OF MCS** | 5000 |
| **INITIAL CELL LAYOUT** | Custom Layout (PIF file) |
| **DEFINED CELL TYPES** | |
| **CELL TYPE "CELL"** | Name of cell type assigned during PIF file generation |
| **CELL TYPE "WALL"** | Pseudo-cell type to prevent cells from sticking to boundaries |
| **CHEMICAL FIELDS** | |
| | No chemical fields were implemented |
| **SELECTED PLUGINS** | |
| **CELLULAR BEHAVIORS:** | |
| ADHESION | Contact |
| **CONSTRAINTS AND FORCES:** | |
| VOLUME | VolumeLocalFlex |
| SURFACE | SurfaceLocalFlex |

| | |
|---|---|
| **CELLULAR PROPERTY TRACKERS** | PixelTracker |
| **CC3D DEFAULT PARAMETERS** | |
| $\lambda_{Volume}$ | 2.0 J/pix$^6$ |
| $\lambda_{Surface}$ | 2.0 J/pix$^4$ |
| $J^{CELL-CELL}$ | 10.0 J/pix$^2$ |
| $J^{CELL-MEDIUM}$ | 10.0 J/pix$^2$ |
| $J^{MEDIUM-MEDIUM}$ | 10.0 J/pix$^2$ |
| **OTHER PARAMETERS** | |
| $J^{CELL-WALL}$ | 500.0 J/pix$^2$ |

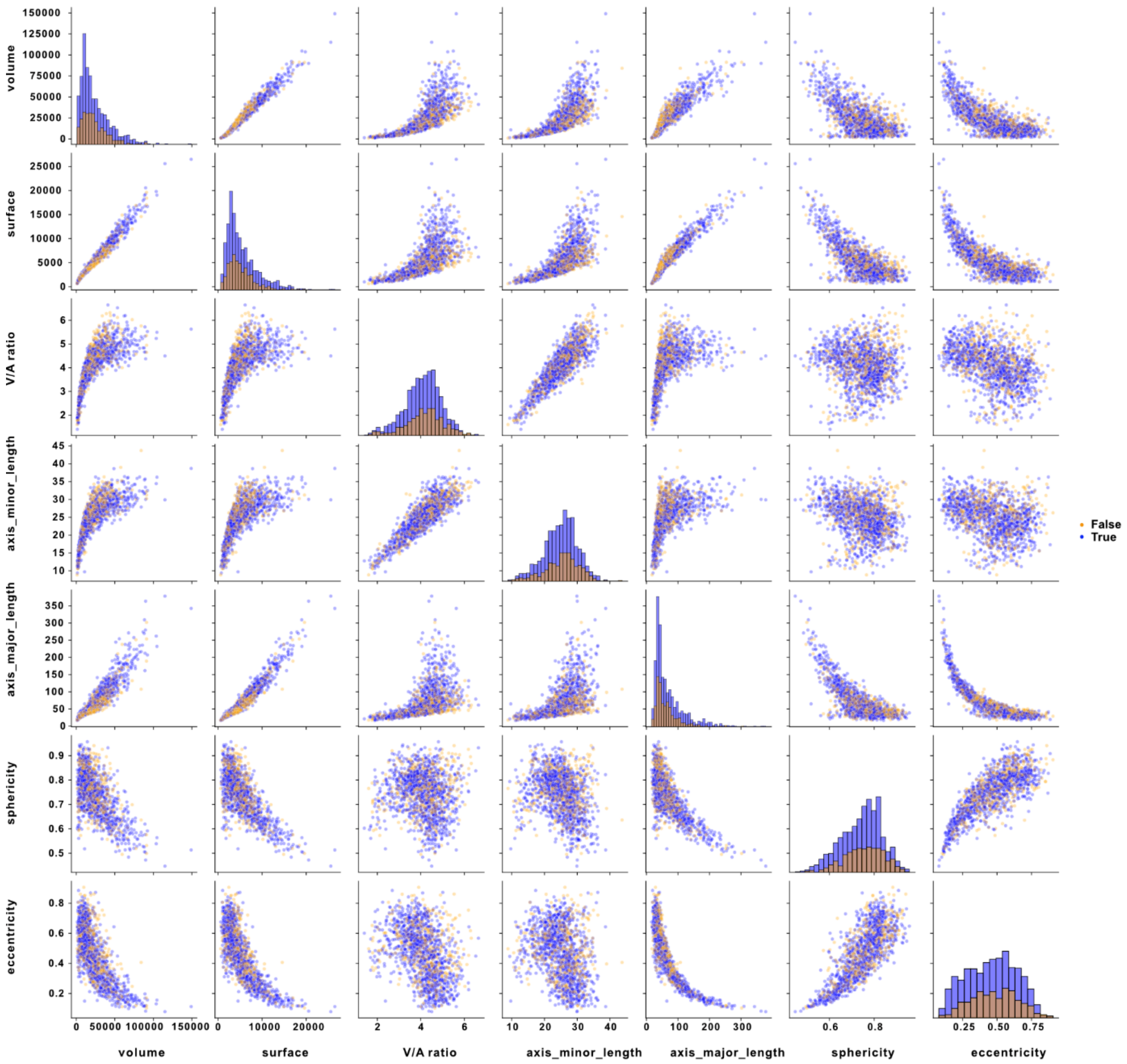


**S 3 Scatter matrix plot of selected single cell morphological features.** Figure displays scatter plots and histograms representing the distribution of various morphological features, highlighting differences between correctly and incorrectly segmented cells. The scatter plots show the relationships between pairs of morphological features, with correctly segmented cells (True) indicated in blue and incorrectly segmented cells (False) in orange. The histograms along the diagonal depict the frequency distributions of individual morphological features for both correctly and incorrectly segmented cells, facilitating the comparison of their distributions.
